# Soil nutrition-dependent dynamics of the root-associated microbiome in paddy rice

**DOI:** 10.1101/2024.09.02.610732

**Authors:** Asahi Adachi, Yuniar Devi Utami, John Jewish Dominguez, Masako Fuji, Sumire Kirita, Shunsuke Imai, Takumi Murakami, Yuichi Hongoh, Rina Shinjo, Takehiro Kamiya, Toru Fujiwara, Kiwamu Minamisawa, Naoaki Ono, Shigehiko Kanaya, Yusuke Saijo

## Abstract

- Plants accommodate diverse microbial communities (microbiomes), which can change dynamically during plant adaptation to varying environmental conditions. However, the direction of these changes and the underlying mechanisms driving them, particularly in crops adapting to the field conditions, remain poorly understood.
- We investigate the root-associated microbiome of rice (*Oryza sativa* L.) using 16S rRNA gene amplicon and metagenome sequencing, across four consecutive cultivation seasons in a high-yield, non-fertilized, and pesticide-free paddy field, compared to a neighboring fertilized and pesticide-treated field.
- Our findings reveal that root microbial community shifts and diverges based on soil fertilization status and plant developmental stages. Notably, nitrogen-fixing bacteria such as *Telmatospirillum, Bradyrhizobium* and *Rhizomicrobium* were over-represented in rice grown in the non-fertilized field, implying that the assembly of these microbes supports rice adaptation to nutrient-deficient environments.
- A machine learning model trained on the microbiome data successfully predicted soil fertilization status, highlighting the potential of root microbiome analysis in forecasting soil nutrition levels. Additionally, we observed significant changes in the root microbiome of *ccamk* mutants, which lack a master regulator of mycorrhizal symbiosis, under laboratory conditions but not in the field, suggesting a condition-dependent role for CCaMK in establishing microbiomes in paddy rice.

## Introduction

Plants host diverse microbial communities (microbiomes), which dynamically change their community structures in response to fluctuating environments. These changes can enhance nutrient acquisition and stress resilience of the host (Fitzpatrick *et al*., 2020). A key aspect of this adaptation is the recruitment and hosting of beneficial microbes in the roots (Lugtenberg & Kamilova, 2009; Berendsen *et al*., 2012). It is well-documented that both abiotic and biotic factors significantly influence the composition and function of root-associated microbiomes. Abiotic factors include climate, soil types, cultivation practices, and the availability of water and nutrients. Biotic factors include host genotypes, root morphology and physiology, and plant developmental stages (Edwards *et al*., 2015, 2018; Zhang *et al*., 2018; Santos-Medellín *et al*., 2021). Accumulating evidence suggests that dynamic alterations and diversification of plant-associated microbiomes play a crucial role in conferring adaptive traits to host plants in various environments (Trivedi *et al*., 2020).

Given the challenges of global food insecurity and climate change, a deeper understanding of plant microbiomes and their potential exploitation is essential to increase crop resilience and productivity for sustainable agriculture (Singh & Trivedi, 2017; Toju *et al*., 2018; Singh *et al*., 2023). Reducing the environmental and labor costs associated with excessive use of chemical fertilizers requires understanding how beneficial microbiomes are established during plant adaptation to nutrient-deficient soils. Rice (*Oryza sativa* L.), a staple crop for half of the world’s population (Edwards *et al*., 2015), has been extensively studied as a model for crop-microbiome research (Kim & Lee, 2020). The rice root microbiome analyses using 16S rRNA amplicon sequencing, conducted on a field-derived soil under greenhouse conditions, have revealed a divergence in microbial communities between the rhizosphere and bulk soil (Breidenbach *et al*., 2016), highlighting the selective hosting of soil-derived bacteria by rice plants (Edwards *et al*., 2015). Among the factors influencing rice root microbiome assembly in paddy fields, fertilization type and degree have been identified as major drivers (Ikeda *et al*., 2014; Aoki *et al*., 2022). Notably, a pot experiment by Sinong *et al*. (2021) using soil from a paddy field left unfertilized for 17 years identified distinct microbial associations related to plant growth promotion in low-nutrient condition. This underscores the need to explore how these findings translate to field conditions, as our understanding of the dynamics and regulatory principles of the root-associated microbiome in response to soil nutrient status is still limited.

Among the mutualistic microbes that assist in plant nutrition, arbuscular mycorrhizal (AM) fungi have garnered significant attention. Under inorganic phosphate (Pi) deficiency, most land plants, including rice, rely on AM symbiosis to acquire Pi from the soil (Singh *et al*., 2023; Inoue *et al*., 2024). This symbiosis is established through the plant’s designated symbiosis (SYM) pathway (Wang *et al*., 2022). The SYM pathway shares key components with the root nodule symbiosis in legumes, including the Ca^2+^/calmodulin-dependent protein kinase (CCaMK), which likely acts as a decoder of Ca^2+^ spiking in response to symbiotic microbes - a critical step in initiating these symbioses (Hayashi *et al*., 2010; Singh & Parniske, 2012). Core SYM pathway genes, including *CCaMK*, are highly conserved in sequence and function across distantly related AM-host plants (Zhu *et al*., 2006), and are also essential for AM symbiosis in rice (Banba *et al*., 2008; Gutjahr *et al*., 2008). However, AM symbiosis may be less effective in inundated paddy fields (Ilag *et al*., 1987). Despite this, studies have shown that the composition of root-associated bacteria changes in *ccamk* rice mutants, which also exhibit reduced plant growth in paddy fields (Ikeda *et al*., 2011). This raises the question about the possible regulation of root microbiomes by CCaMK, particularly in relation to soil nutrition status.

In this study, we utilized a paddy field that has not been treated with chemical fertilizers or pesticides for approximately 70 years yet has consistently yielded at least 60% of the crop output compared to conventionally fertilized fields in the same region. We hypothesized that a root-associated microbiome adapted to nutrient-limiting soil conditions is established in this specific environment. To investigate this, we compared the community structure of the root-associated microbiome, focusing on bacteria and archaea, of rice grown in this long-term, non-fertilized paddy field to that in a neighboring, conventionally fertilized field. We assembled 16S rRNA sequence profiles from the root endosphere, a possible niche for the “core microbiome” (Edwards *et al*., 2015), sampled throughout the four consecutive growing seasons from 2018 to 2021. Using these data, we explored the feasibility of building a random forest classification model to predict soil fertilization status based on the relative abundance of specific microbes. Additionally, we examined the possible role of CCaMK in the establishment of root microbiota by using *ccamk* mutant plants.

## Materials and Methods

### Rice materials, fields and sampling procedures

Rice roots were sampled from fertilized and non-fertilized fields in Uji, Kyoto Prefecture, Japan (approximately at 34.9°N latitude and 135.8°E longitude). The non-fertilized field, whose soil has not received fertilizers since 1951, has been noted for high productivity (about 300 kg/10 a) (Kobayashi, 2015). In contrast, the fertilized field has been supplied with 20-25 kg/ 10 a of organic fertilizers every year (Kyoto Yamashiro Organic, Kyoto, Japan). Both fields are adjacent but have separate irrigation systems. We cultivated three Japonica varieties of rice (*Oryza sativa*): Nipponbare, Hinohikari, Kinmaze, and *ccamk* mutants (in the Nipponbare background with one Tos17 insertion) (Hirochika, 2001; Miyao *et al*., 2003), in these two fields. Rice roots were sampled every two or three weeks from the vegetation phase to harvest over four growing seasons from 2018 to 2021. Root samples were washed, cooled, and transported to the laboratory, where they were surface sterilized with soap, 70% ethanol for 1 min, and 2% hypochlorous acid for 1 min, and then washed with sterile water, before their storage at −80°C.

Additionally, pot experiments with Nipponbare WT and *ccamk* mutants were conducted in the greenhouse. We used a soil mixture consisted of the Ogura non-fertilized soil, Akadama soil (extra small grain; Plantation Iwamoto, Ibaraki, Japan), and Kanuma soil (fine grain; Akagi Engei, Gunma, Japan) in a 4:5:1 mass ratio. Rice roots were sampled at 6, 10, 12, and 17 weeks after germination. Urea (KOMERI Co., Ltd., Niigata, Japan) was used as a nitrogen source, and superphosphate (SUN and HOPE Co., Ltd., Fukuoka, Japan; KOMERI Co., Ltd., Niigata, Japan) provided phosphate. Sampling followed the same procedure as in the field.

### Element analyses

The nitrogen and phosphorus concentrations in the rice shoots were determined using JM3000CN analyzer (J-Science Lab Co., Ltd., Kyoto, Japan) and a colorimetric method for hydrochloric acid-soluble fractions (Murphy & Riley, 1962). Element concentrations in the flag leaves of rice were determined by inductively-coupled plasma-mass spectrometry (ICP-MS) as described by Tanaka *et al*., 2018. The properties and nutrient availability of the field soils were analyzed with inductively coupled plasma optical emission spectrometry (ICP-OES) at Yanmar Holdings Co., Ltd. (Tokyo, Japan; https://www.yanmar.com/global/agri/soil_solution/).

### DNA extraction and 16S rRNA gene amplicon sequencing

Root samples were ground using Mixer Mill (Verder Scientific Co., Ltd., Tokyo, Japan), and total genomic DNA was extracted with NucleoSpin® Soil (MACHEREY-NAGEL GmbH & Co. KG, Germany) according to the manufacturer’s instruction. V4 region of the 16S rRNA gene was amplified by touchdown PCR using KOD FX Neo (TOYOBO Co., Ltd., Osaka, Japan) and Illumina barcoded 515 forward (TCGTCGGCAGCGTCAGATGTGTATAAGAGACAG-GTGCCAGCMGCCGCGGT AA) and 806 reverse (GTCTCGTGGGCTCGGAGATGTGTATAAGAGACAG-GGACTACHVGGGTWTCT AAT) primers according to Edwards *et al*. (2015). Bands of the target DNA were excised from 1% agarose gel electrophoresis and purified using Fast Gene Gel/PCR extraction kit (Nippon Genetics Co., Ltd., Tokyo, Japan) following the kit procedure. In the following steps, amplicon libraries were prepared using Nextera XT Indices (Illumina, Inc., California, USA) and paired-end sequencing was performed on the Illumina MiSeq platform using the MiSeq Reagent Kit v3.

### Sequence data processing

Raw sequence data were demultiplexed using Claident software (Tanabe & Toju, 2013). Amplicon sequence variants (ASVs) were identified with the DADA2 (Callahan *et al*., 2016) using the dada2 package in R. The parameters of filterAndTrim function were set to the following: truncQ = 2, truncLen = 220 and 150, trimLeft = 19 and 20, maxN = 0, and maxEE = 2 and 2 for the forward and reverse reads, respectively. Chimeric sequences were identified and removed with default settings. Taxonomy for representative ASV sequences was assigned against the Silva 138.1 database (Quast *et al*., 2013; Yilmaz *et al*., 2014) with assignTaxonomy function. ASVs classified as *Eukaryota* or not assigned to any domain, as well as those from mitochondria and chloroplasts, were excluded.

### Statistical analyses

To assess and compare the alpha diversity (within-sample diversity), we used the Shannon index (Shannon, 1948) calculated for each microbiome sample based on relative ASV abundance with the ‘diversity’ function of the vegan package (Dixon, 2003) in R. We performed Welch’s t-test to assess the effects of fertilization, with *p*-values corrected using the Bonferroni method. We conducted Jonckheere-Terpstra test (Jonckheere, 1954) using clinfun package in R, with 1000 permutations for the reference distribution. Additionally, we calculated the Bray-Curtis dissimilarities between samples using the vegan package in R, and used the Bray-Curtis dissimilarities for principal coordinates analysis (PCoA) and permutational multivariate analysis of variance (PERMANOVA) (Anderson, 2001) to reveal microbiome structures. PERMANOVA was performed using the ‘adonis2’ function in the vegan package, with 999 permutations. All statistical analyses were performed in R, and plots were created using the ggplot2 package (Wickham, 2016).

To identify microbes more abundant in non-fertilized fields compared to fertilized fields, we conducted differential abundance testing with ALDEx2 (Fernandes *et al*., 2014). ALDEx2 converts count values to probabilities through Monte Carlo sampling from the Dirichlet distribution with a uniform prior, applies a center log-ratio (CLR) transformation, and then performs Welch’s t-tests. *p*-values were adjusted using the false discovery rate (FDR) approach of Benjamini-Hochberg. This analysis used the relative abundance data of Nipponbare root microbiomes across four growing seasons (2018-2021) with ALDEx2 executed using the ‘aldex’ function in the ALDEx2 package in R.

### Random forest classification

We developed a random forest (RF) classifier (Breiman, 2001) to predict soil fertilization states (i.e., fertilized or non-fertilized) based on microbiome profiles. Given the high-dimensional nature of microbiome data, where the number of features often exceeds the number of samples (Asnicar *et al*., 2024), we aggregated features based on taxonomical information to manage dimensionality (Duvallet *et al*., 2017). Specifically, we collapsed ASVs to the genus level and used genus-level relative abundances as input features. ASVs not assigned to any genus were excluded.

We trained the RF model using genus-level microbiome profiles of Nipponbare from three growing seasons (2018-2020). The ‘mtry’ parameter of the RF was optimized using 5-fold cross-validation on the training data. To accurately assess model performance, we used independent test data from different years and rice genotypes: Nipponbare (2021), Hinohikari (2019-2020), Kinmaze (2018), and *ccamk* mutants (2019-2021). The training of RF and performance evaluation was conducted using the caret package (Kuhn, 2008) in R.

To investigate how classification accuracy varied with the rice growth stage, we analyzed weekly changes in accuracy. For this, we used Nipponbare samples (2018-2021), training the model on data from three years and testing it on the remaining year. Accuracy for each week was computed as the average of accuracies over five weeks, including two weeks before and after the target week.

### Metagenome sequencing and analyses

DNA samples of the Nipponbare cultivar were collected at three time points during the 2019 season - 8, 15, and 20 weeks - corresponding to the vegetative, reproductive, and seed maturation phases, respectively. Metagenomic sequencing was conducted at Takara Bio Inc. (Shiga, Japan). Briefly, DNA was fragmented using Covaris sonicator (Covaris, Massachusetts, USA) and cleaned with AMPure XP beads (Beckman Coulter, California, USA). Libraries were prepared with SMARTer ThruPLEX DNA-Seq Kit (Takara Bio Inc., Shiga Japan) and sequenced as 150bp pair-end reads on NovaSeq 6000 platform (Illumina, Inc., California, USA) using NovaSeq 6000 S4 Reagent Kit.

Sequence reads were processed for adapter and quality trimming using the cutadapt v1.8 (Martin, 2011) and PRINSEQ v0.20.4 programs (Schmieder & Edwards, 2011). Clean reads were aligned against the *O. sativa* genome (accession: NW_015379174.1) to remove host contamination using bowtie (Langmead *et al*., 2009). Qualified reads were assembled using MEGAHIT (Li *et al*., 2016) to construct contigs. Due to low read mapping rates to contigs in some samples, functional and phylogenetic annotations were performed directly from raw read fragments (R1 reads with a minimum length of 100 bp), bypassing assembly.

Functional annotation was conducted by searching the KEGG prokaryote protein database with MMSeqs2 (Steinegger & Söding, 2018) in blastx mode, using parameters: E-value 1e-5, identity ≥ 40%, and query coverage ≥ 50%. Top hits were phylogenetically classified using the NCBI nr database in MMSeqs2 easy-taxonomy mode.

Functional annotation results were used for further analyses. Metabolic pathway completeness was assessed using KEGGDecoder (Graham *et al*., 2018). Principal component analysis (PCA) based on Bray-Curtis dissimilarities from KEGG Orthology (KO) relative abundance was visualized with ggplot2. KOs with significant differences between fertilized and non-fertilized samples were identified using Boruta v7.0.0 (Kursa & Rudnicki, 2010) and visualized with ggpubr.

## Results

### Microbiome profiling and diversity analyses

We examined the dynamics of the rice root-associated microbiome and its dependence on soil nutrition (fertilization) in neighboring non-fertilized and fertilized paddy fields, each with separate water systems. In this setup, geographical and environmental factors other than soil conditions (as detailed in Table S1) are considered negligible in influencing the developmental transitions and physiology of the rice plants, allowing for a focused interpretation of the results. The non-fertilized field has maintained a high yield despite approximately 70 years without fertilization, prompting our exploration of a root-associated microbiome that may support rice adaptation to nutrient-deficient paddy soils. To examine this, we collected root tissues every 2-3 weeks throughout the rice plant lifecycle, using three Japonica varieties - Nipponbare, Hinohikari, and Kinmaze - over one to four growing seasons from 2018 to 2021. These efforts resulted in a total of 300 root samples, as outlined in Table S2.

To investigate the bacterial and archaeal communities inhabiting the rice root endosphere, we sequenced the V4 region of the 16S rRNA gene from DNA extracted from surface-sterilized roots. Our analyses identified a total of 33,340 amplicon sequence variants (ASVs) across all root samples. These ASVs were classified into two domains, spanning 69 phyla, 155 classes, 348 orders, 454 families, and 1036 genera. Consistent with previous studies (Edwards *et al*., 2015; Xu *et al*., 2020), the most abundant phylum was *Proteobacteria*, representing 63.9% of the relative abundance. This was followed by *Actinobacteriota* (8.1%), *Firmicutes* (7.6%), *Bacteroidota* (4.6%), and *Myxococcota* (2.7%).

We assessed the alpha diversity (within-sample diversity) of the root microbiome samples from the three rice genotypes under the two fertilization conditions by calculating the Shannon index. As shown in Figure 1a, the Shannon index varied depending on both the soil fertilization conditions and the rice cultivars. Across all cultivars, a trend of a higher Shannon index values was observed in the fertilized field compared to the non-fertilized field, consistent with the findings of Ikeda *et al*. (2014) for nitrogen-rich soil. However, the differences in Shannon index between the fertilized and non-fertilized fields were not statistically significant for all cultivars except Hinohikari (adjusted *p* = 0.011). Moreover, the Shannon index increased significantly as the rice plants developed, indicating a rise in species diversity from the young seedling stage to maturity, regardless of genotype and soil fertilization condition (Jonckheere-Terpstra test, *p* = 0.001; Figure 1b).

**Figure 1.**
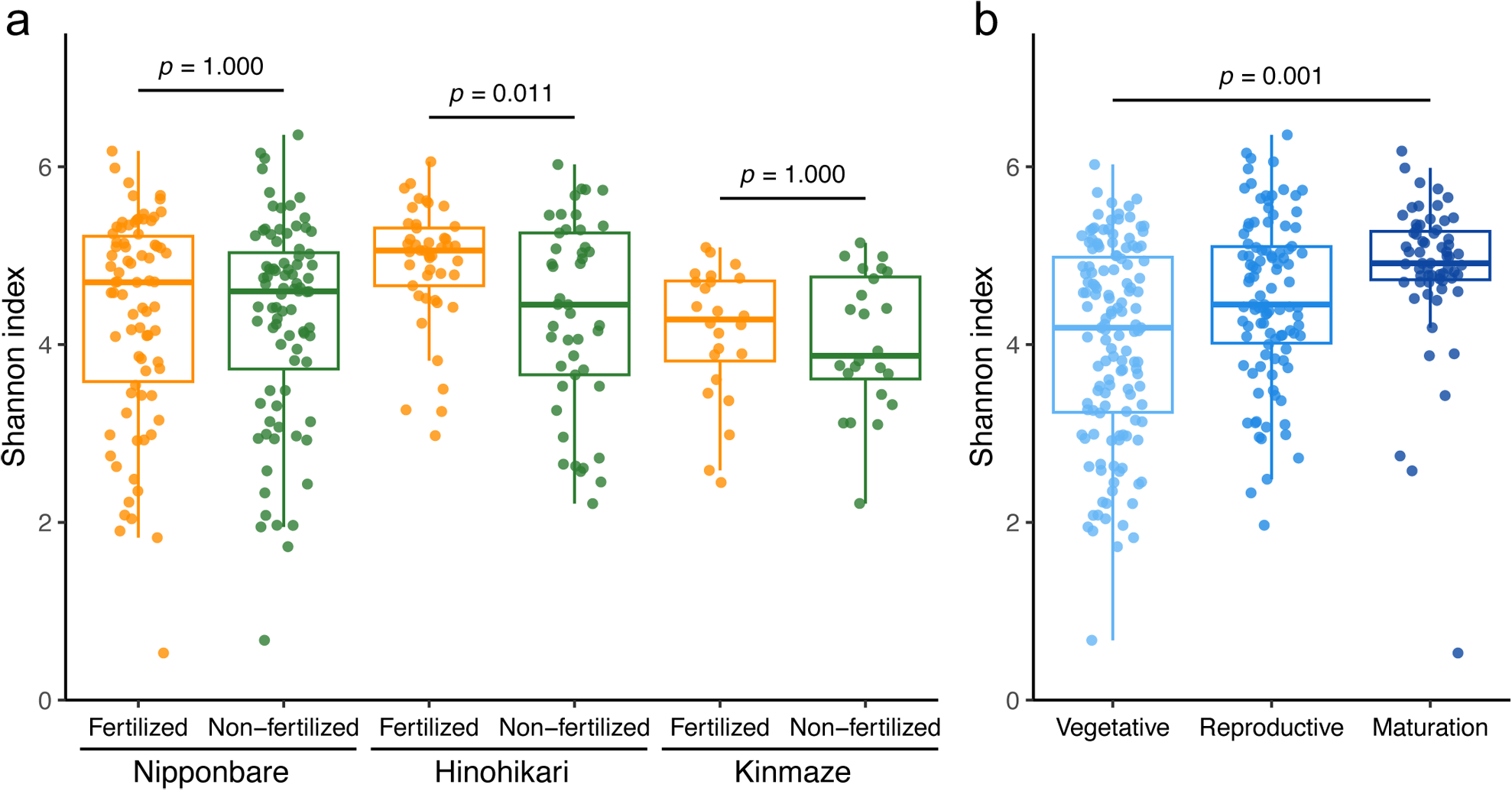
Alpha-diversity of the root endosphere microbiome of field-grown rice sampled from 2018-2021. (a) Shannon index (within-sample diversity) of the root-associated microbiomes of three rice cultivars. Colors indicate different fertilization conditions. Significant differences in microbiome diversity due to soil fertilization were observed in Hinohikari but not in the other two varieties. To assess these differences, we used two-sided unpaired Welch’s t-tests and applied Bonferroni correction to adjust the *p*-values for multiple comparisons. (b) The Shannon index of the root-associated microbiomes is presented, with colors representing physiological stages of rice plants. Samples are categorized into three physiological stages based on weeks after germination: vegetative (≤ 10 weeks), reproductive (≤ 16 weeks), and seed maturation (>16 weeks). The Shannon index increased with the developmental stages of the rice plants, as indicated by the Jonckheere-Terpstra test (*p* = 0.001).

Next, we evaluated the beta diversity (between-sample diversity) of the root-associated microbiomes by calculating the Bray-Curtis dissimilarities. To identify the driving forces behind the divergence and convergence of microbial community structures, we employed principal coordinates analysis (PCoA) (Gower, 1966) along with permutational multivariate analysis of variance (PERMANOVA) (Anderson, 2001) on the Bray-Curtis dissimilarities. The PCoA plots (Figure 2a) illustrate variations in ASV composition among the microbiome samples, showing that the community structures of the root endosphere microbiomes were distinct between fertilized and non-fertilized paddy fields across all cultivars and throughout the growing seasons. The PERMANOVA results confirmed the significant divergence between the fertilized and non-fertilized root microbiomes (Table S3). Both soil fertilization conditions and plant genotypes were significant factors (*p*-values were both 0.001), but the effect of soil fertilization was more substantial compared to plant genotypes (Table S3). These findings align with previous studies that soil type is a major source of variation in the microbial communities of the rice root endosphere (Edwards *et al*., 2015, 2018).

**Figure 2.**
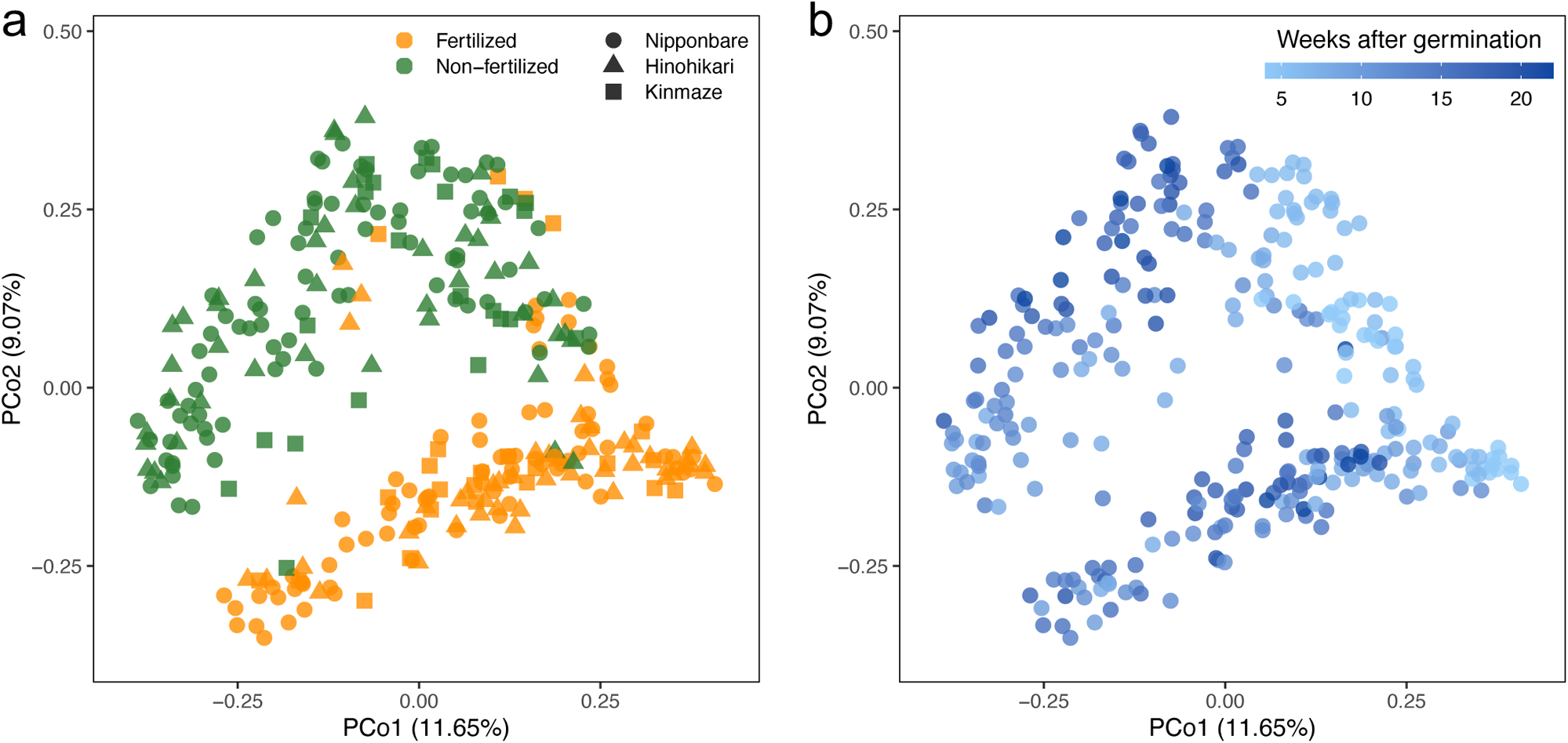
The structure of the root endosphere microbiome varies with soil fertilization conditions and the host plant’s developmental stages. (a) The principal coordinate analysis (PCoA) plot of Bray-Curtis dissimilarities shows microbiome samples distinguished by soil fertilization conditions (color) and rice genotypes (shape). (b) The same samples are colored by plant developmental stages post-germination. Soil fertilization conditions separated the root-associated microbiomes into two distinct groups, each showing significant shifts corresponding to the rice developmental stages.

We then examined how the developmental stages of rice plants affect the root microbiomes. The PCoA plots suggest that microbiome structures undergo noticeable shifts as the plants progress through different developmental stages during the growing seasons, both in the vegetative and reproductive stages, and in both fertilized and non-fertilized fields (Figure 2b). This plant age-dependent shift in the root microbiome aligns with the findings of Zhang *et al*. (2018) and Edwards *et al*. (2018). The results also suggest a divergence in the direction of these shifts between the two fields, implying that the development and diversification of the root endosphere microbiome over the rice plant’s lifecycle may be linked to plant adaptation to soils with different nutrition states.

Our data are in good agreement with Edwards *et al*. (2018) regarding the effects of soil conditions and plant age on the root endosphere microbiomes. In our study, the continuous diversification of the root microbiomes along the progression of plant growth stage was observed in both fields over four growing seasons (Figure 2b). This appears to be in marked contrast to the results of Edwards *et al*. (2018) that the endosphere microbiomes of rice planted in two geographically separate sites, albeit quite dissimilar in the beginning, reach increased levels of similarity by the end of the growing season. However, a common conclusion from both studies might be that the root endosphere microbiomes converge toward similar community structures during the reproductive phase. These structures are likely influenced by the soil nutrient status, even when the initial colonizers from the bulk soil are different.

### Identification of microbes influenced by soil fertilization conditions

We next investigated the specific bacterial groups that contributed significantly to the observed variations in the microbiome structures of the rice root endosphere grown in field conditions. When analyzing the relative phylum abundance, we did not observe any discernible common patterns across the three cultivars between fertilized and non-fertilized fields, nor did we see consistent changes with plant age (Figure 3). However, in Hinohikari and Kinmaze, but not in Nipponbare, the ratio of *Actinobacteriota* increased in the fertilized field compared to the non-fertilized field (Figure 3). We then performed differential abundance analyses with ALDEx2, which identified ASVs that were significantly more abundant in either fertilized or non-fertilized fields. Figure 4 presents the effect size of these ASVs across different growing seasons and rice genotypes. These differential ASVs were identified through the Benjamini–Hochberg correction of the *p*-value (adjusted *p* < 0.05).

**Figure 3.**
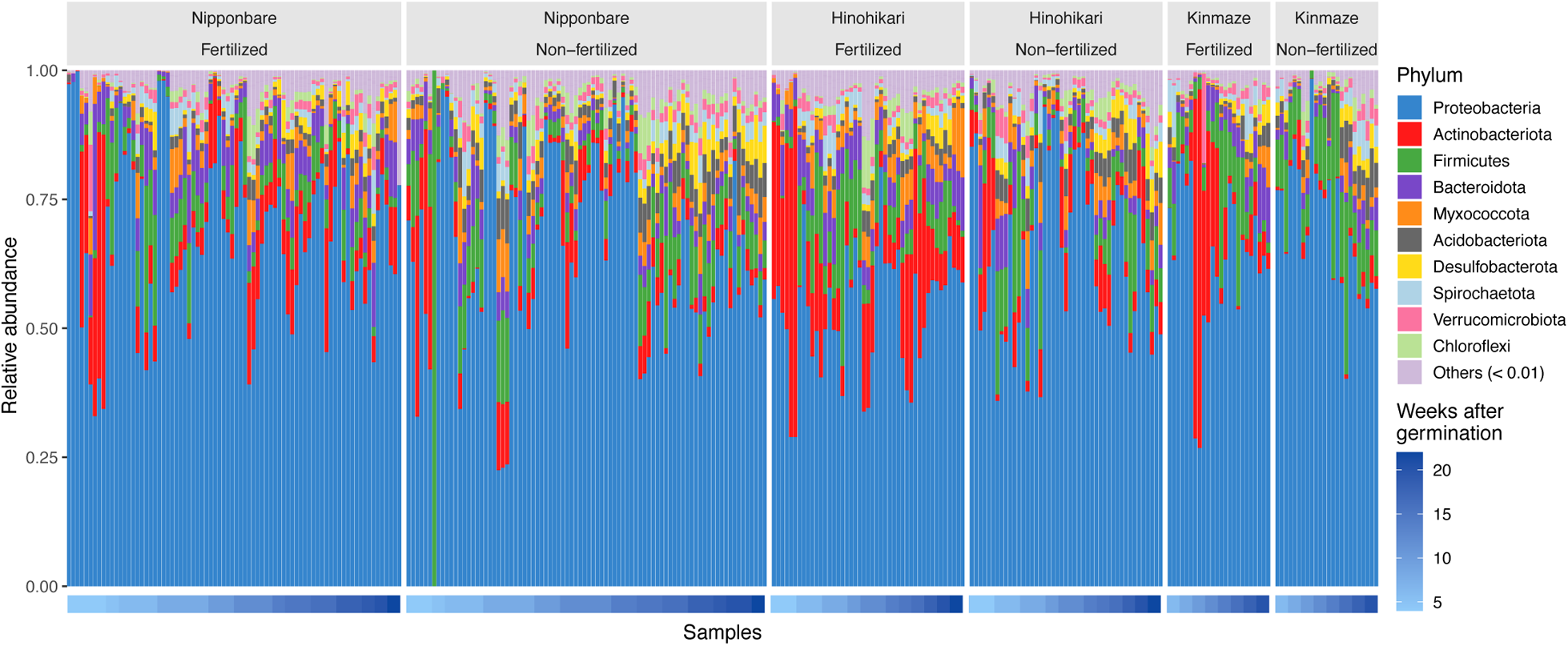
Phylum-level relative abundance of ASVs in the rice root microbiome for 300 field samples. The samples are arranged in a time series, sorted by weeks post-germination at harvest. Each phylum is represented by a different color, as indicated in the legend on the right.

**Figure 4.**
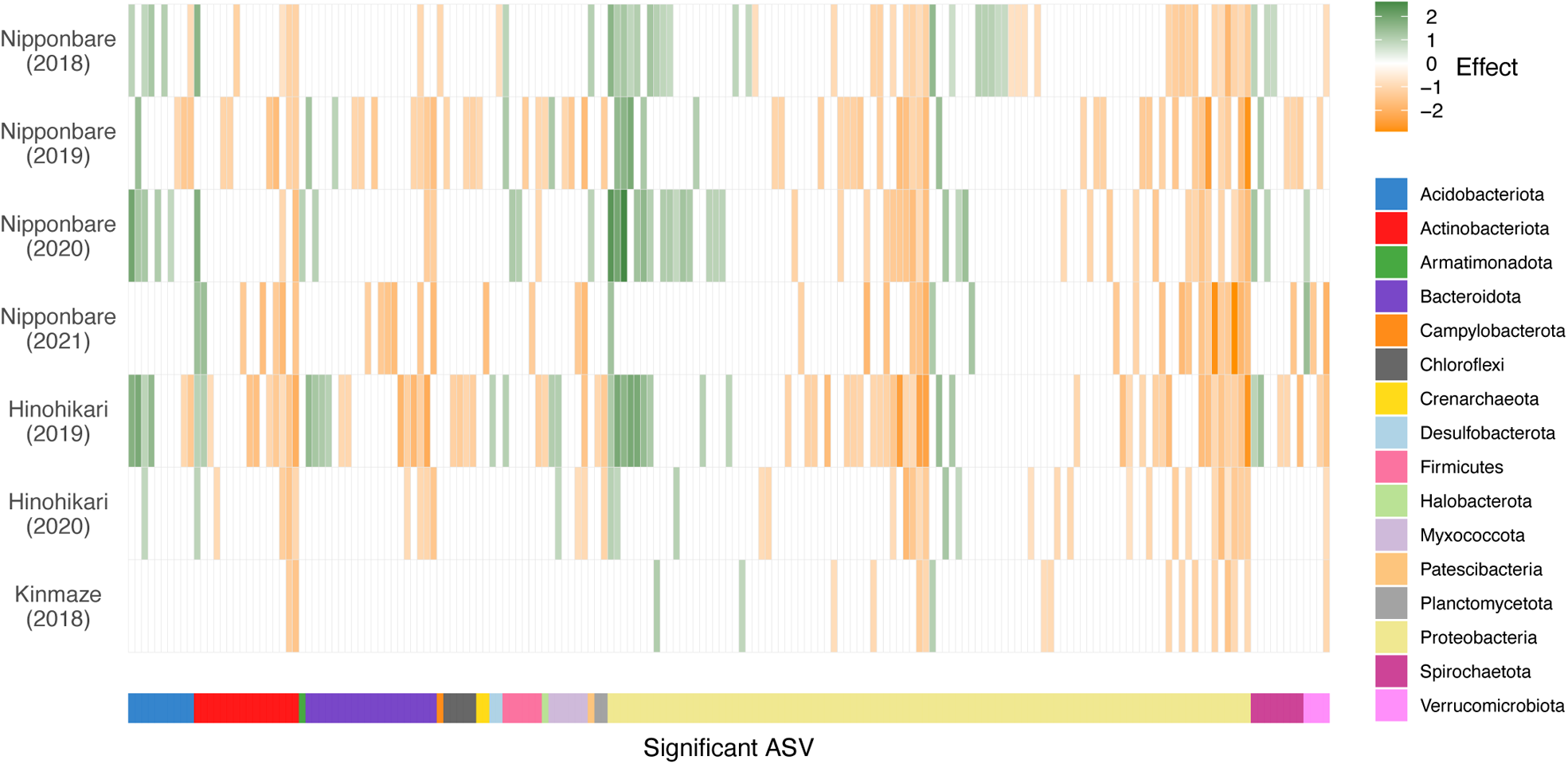
Differential abundance of ASVs in fertilized and non-fertilized fields, revealed by ALDEx2 analysis. All significant amplicon sequence variants (ASVs) that were over-represented in either fertilized or non-fertilized fields, in at least one genotype-year combination, were identified using ALDEx2. These ASVs were filtered based on p-values after applying the Benjamini–Hochberg correction (adjusted *p* < 0.05). The effect size of each ASV, relative to the non-fertilized field, is depicted using a color gradient. ASVs significantly more abundant in the non-fertilized field are shown in green, while those more abundant in the fertilized field are shown in orange. The phyla corresponding to these ASVs (shown in the bottom) are represented by different colors, as indicated in the legend on the right.

The analysis revealed that 183 ASVs exhibited significant differences in relative abundance between fertilized and non-fertilized fields, in at least one growing season and genotype combination. Of these, 61 ASVs were over-represented in rice roots from the non-fertilized field, whereas 122 ASVs were over-represented in the fertilized field. Notably, most phyla included ASVs from both groups, except for *Chloroflexi* and some phyla represented by only 1-2 ASVs, suggesting diversification within different phyla based on soil fertilization preference (Figure 4). No ASVs displayed stochastic distribution patterns, such as being over-represented in different growing seasons or genotypes between the two fields, indicating a high level of consistency in bacterial/archaeal preference for fertilization conditions at the ASV level. This consistency suggests that these ASVs could serve as reliable diagnostic markers for soil fertilization status.

ASVs over-represented in the non-fertilized field belonged to the bacterial groups known for nitrogen-fixing species, such as the genus *Azospirillum* (Baldani *et al*., 2015b), the genus *Bradyrhizobium* (Kuykendall, 2015), and the genus *Herbaspirillum* (Baldani *et al*., 2015a) (Table S4). This finding supports our hypothesis that a root microbiome adaptive to aiding plant nutrition acquisition is established in high-yield, non-fertilized fields.

### Environmental context-dependent role of CCaMK in regulating root endosphere microbiomes

We next explored a possible role for CCaMK in regulating plant growth and the root-associated microbiome under inundated paddy field soil conditions. To this end, we compared the root endosphere microbiomes between wild-type Nipponbare and *ccamk* mutant rice grown in both fertilized and non-fertilized fields (Table S5), as well as in pots in a greenhouse. We analyzed both alpha and beta diversity between the wild-type (WT) and *ccamk*.

Figures 5a and 5b display the Shannon indices of root-associated microbiomes from rice grown under different fertilization conditions in the field and greenhouse, respectively. An unpaired Welch’s t-test was used to compare the effects of *CCaMK* disruption and soil fertilization, with *p*-values adjusted for multiple comparisons using the Bonferroni correction. In both fertilized and non-fertilized fields, the Shannon index of the root microbiome did not significantly differ between the WT and *ccamk* plants, a conclusion also reached for plants grown in the greenhouse. However, in *ccamk* plants, the Shannon index of the root microbiome differed significantly between the fertilized and non-fertilized fields (adjusted *p* < 0.001), a pattern not observed in WT plants. Conversely, no significant difference in the Shannon index depending on fertilization status was found in *ccamk* plants grown in the greenhouse, while a significant difference was observed in WT plants (adjusted *p* = 0.019). Thus, no consistent trend in the alpha diversity was obtained between the field and greenhouse conditions.

**Figure 5.**
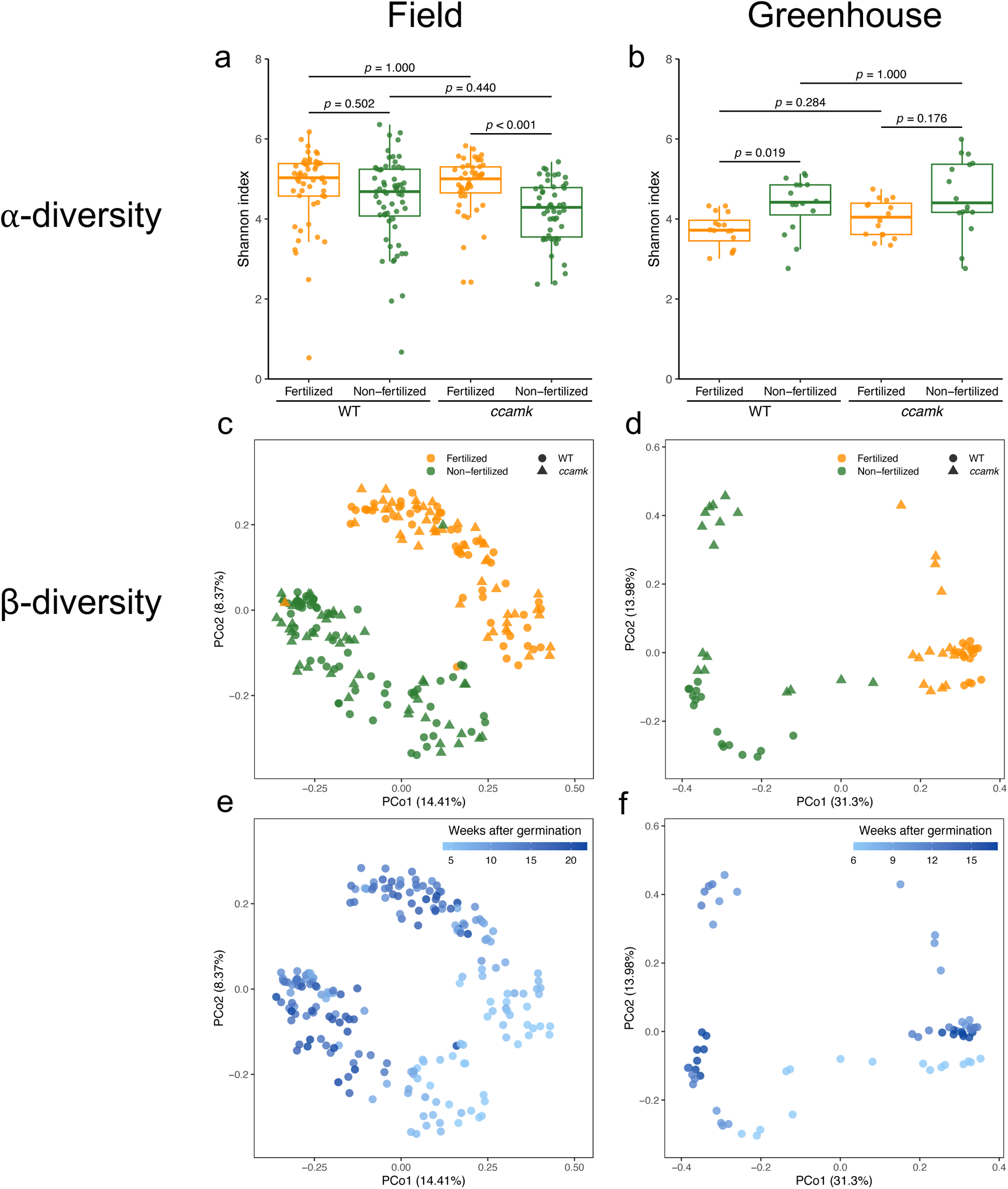
Alpha and beta diversity of root endosphere microbiomes of the wild-type and *ccamk* plants grown in the field and greenhouse. Samples were collected from the field (2019-2021) (a, c, and e) and the greenhouse (2020) (b, d, and f). The alpha diversity (Shannon index) of root-associated microbiomes was analyzed (a and b). Two-sided unpaired Welch’s t-tests were performed to evaluate the impact of soil fertilization, with *p*-values adjusted for multiple comparisons using the Bonferroni correction. The beta diversity of root microbiomes showed significant differences between soil fertilization conditions (c and d) and across plant developmental stages (e and f). In the greenhouse, beta diversity also varied significantly depending on *CCaMK* (d).

Figures 5c and 5d illustrate that the beta diversity of the root microbiomes varied significantly under different soil fertilization conditions, both in the field and in the greenhouse. In line with previous findings, plant age also significantly influenced the beta-diversity of microbiomes in both environments (Figures 5e and 5f). In field-grown rice, a trend of lower biomass was observed in *ccamk*, particularly in the non-fertilized field, similar to the results reported by Ikeda *et al*. (2011) (Figure S1). However, *CCaMK* disruption did not significantly impact the beta-diversity of the root microbiomes in the field (Table S6). On the other hand, while the soil fertilization condition was the dominant factor affecting beta diversity (Table S7), the presence of the *CCaMK* gene also significantly influenced beta diversity of root microbiomes in rice grown in the greenhouse. These findings reinforce that soil fertilization condition and plant developmental stage play critical roles in shaping beta diversity of the rice root microbiome. However, the influence of the *CCaMK* gene on beta diversity appears to be dependent on environmental context.

### Predicting soil nutrition status using microbiome data through a machine learning model

Given the strong associations observed between specific microbes and fertilization conditions, we next aimed to develop a machine learning model to predict soil fertilization status based on the relative abundance of the root microbiomes. We trained a random forest (RF) classification model to distinguish whether a plot was fertilized or not, using 16S rRNA gene amplicon sequencing data. The model utilized genus-level aggregated relative abundance of each ASV as features.

Initially, we trained the RF model using data from the Nipponbare cultivar collected over three growing seasons (2018-2020). We then assessed the model’s robustness using multiple test datasets. The classification accuracies across different testing datasets are detailed in Figure 6a. When applied to the Nipponbare data in 2021, the model achieved a classification accuracy of 0.900, highlighting its robustness against year-to-year variations. We further evaluated the model’s robustness across different rice genotypes using the data from Hinohikari (2019-2020), Kinmaze (2018), and *ccamk* mutants in the Nipponbare background (2019-2021). The model, trained on Nipponbare (2018-2020) data, achieved classification accuracies of 0.944, 0.875, and 0.949 for Hinohikari, Kinmaze, and *ccamk* mutants, respectively (Figure 6a). These results demonstrate that our RF model effectively predicts soil nutrition status across different growing seasons and rice genotypes in the study fields. The findings suggest that specific microbial subsets in the root endosphere consistently follow distinct relative abundance patterns across the plant life cycle, depending on the soil fertilization status.

**Figure 6.**
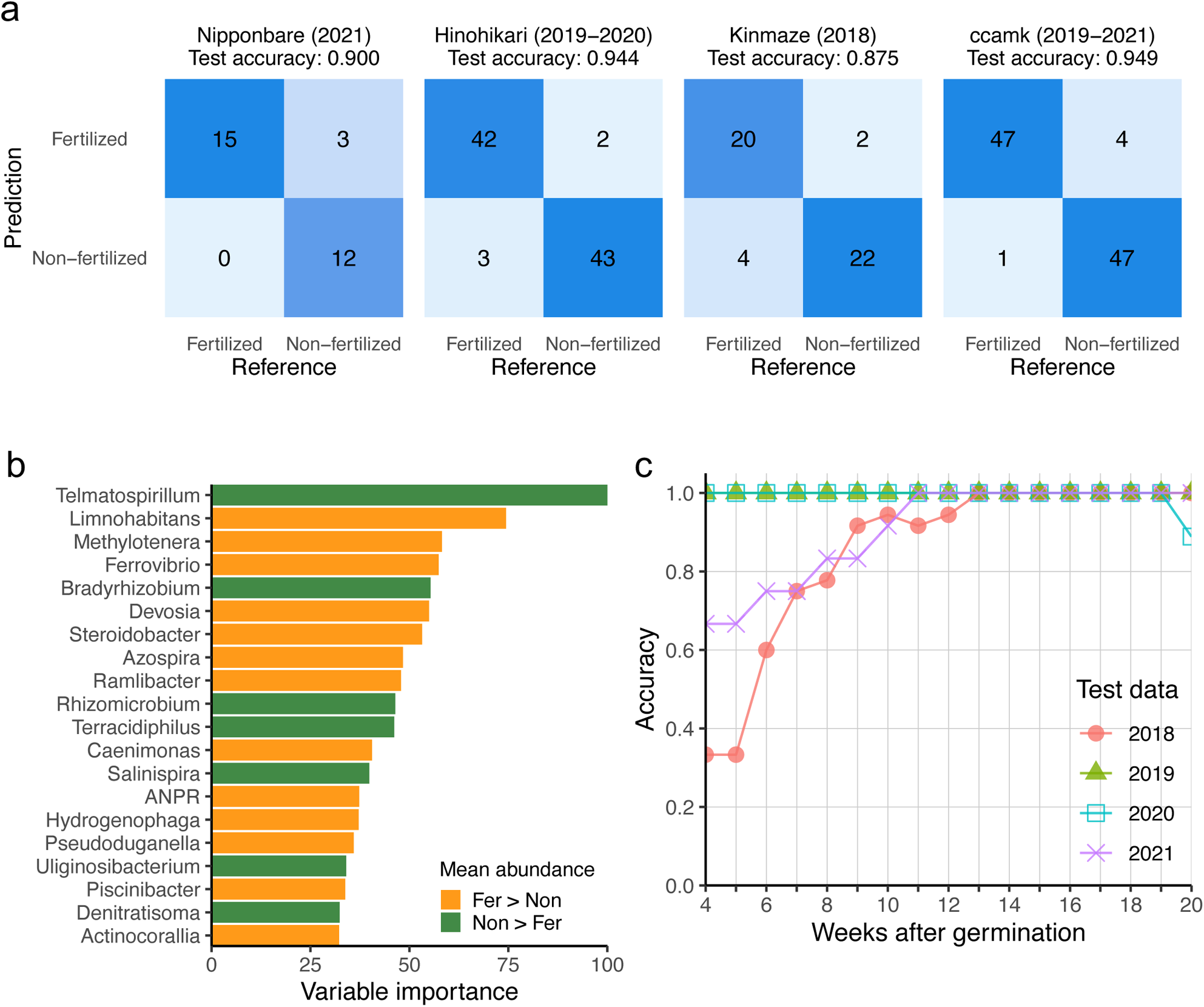
Building a random forest classification model to predict soil fertilization status on the root microbiome data. a) The robustness of the random forest (RF) classification model, trained on datasets from three growing seasons (2018-2020) of Nipponbare, is demonstrated through confusion matrices for four test datasets: Nipponbare (2021), Hinohikari (2019-2020), Kinmaze (2018), and *ccamk* mutants (2019-2021). b) The top 20 features of the RF classification model are ranked by their importance. Each feature’s abundance relative to soil fertilization conditions is indicated: orange (fertilized field > non-fertilized field), green (non-fertilized field > fertilized field). ANPR refers to a group of genera: Allorhizobium, Neorhizobium, Pararhizobium, and Rhizobium. (c) The classification accuracy of the RF model is evaluated using Nipponbare data from 2018-2021. In this evaluation, each year was used as a test dataset, while the other three years were used for training. This method evaluates the model’s performance and accuracy across different sampling time points.

We then visualized the variable importance of the RF model features to identify the genera that contributed most to the accurate prediction of soil fertilization status. The top 20 genera by variable importance are presented in Figure 6b. We examined the abundance of these genera in the bulk soil and found that some, such as *Limnohabitans*, *Azospira* and *Rhizomicrobium,* may have originated from the native soil microbiota and become enriched in the root endosphere (Figure S2c). Notably, *Telmatospirillum*, the top genus enriched in the non-fertilized field, did not show differential abundance in the bulk soils of fertilized versus non-fertilized fields (Figure S2c), suggesting that it may be selectively enriched in the rice root endosphere in response to soil conditions. Isolates from the genus *Telmatospirillum* have been reported to possess nitrogen-fixing (Sizova *et al*., 2007) and iron-reduing (Gagen *et al*., 2019) capabilities. Additionally, the enrichment of other nitrogen-fixing genera, such as *Rhizomicrobium* (Li *et al*., 2021; Wang *et al*., 2023) and *Bradyrhizobium* (Itakura *et al*., 2009), in the non-fertilized field further supports our hypothesis of a plant microbiome that adapts to the soil nutrient status.

We also assessed whether prediction accuracy varied with the developmental stages of rice. RF models were trained on Nipponbare data from three of the four growing seasons, and then tested for accuracy on Nipponbare data from the remaining year. Figure 6c illustrates the accuracy scores for the four training-testing dataset pairs, where each week’s accuracy represents the average of the accuracies over five weeks (including the two weeks before and after the target week). The accuracy was initially low during the first 4-7 weeks after germination, but peaked around 13-19 weeks after germination, coinciding with the vegetative-to-reproductive phase transition, before declining toward 20 weeks after germination. This indicates that root microbiome data collected during the 13-19 weeks post-germination is most effective for distinguishing between fertilized and non-fertilized conditions using the RF model. In contrast, root microbiome data from the initial 5-6 weeks after germination (1-2 weeks after seedling transfer to paddy fields) and the final weeks before harvest showed lower predicting power, suggesting that fertilization-discriminant genera’s relative abundance patterns are less pronounced in the beginning and end of the growing season.

Overall, our results suggest that the endosphere microbiomes established around the vegetative-to-reproductive phase transition significantly influence the accuracy of the soil fertilization condition prediction models. While robust models have been developed to predict soil health based on soil microbiomes (Wilhelm *et al*., 2022; Dai *et al*., 2023), our findings indicate that soil properties or fertilization states can be indirectly forecasted by using the root endosphere microbiome as an indicator.

### Functional annotation of the root endosphere microbiome using metagenomic sequencing analyses

Metagenomic sequencing analyses were also carried out on the Nipponbare root samples at three time points during the 2019 growing season, which represent vegetative, reproductive, and seed maturation phases. Functional annotation was conducted using KEGG resulting to the collection of 9,304 KEGG Orthologues (KOs). The detected KOs can be assigned into 169 types of metabolic pathways based on KEGGDecoder. The data at young stages have lower completeness of metabolic pathways than older ones, both in the fertilized and non-fertilized fields. However, it is conceivable that incompleteness of some metabolic pathways, due to the lack of some of the key genes, was rather attributable to their low abundance below the detection levels in our sequencing analyses than their absence. Nevertheless, the low read counts likely reflect the low number of the microbial cells possessing these genes in their genomes. Therefore, we used the metabolic pathway completeness in the metagenome analyses as a proxy for the extent to which the microbes are represented in the samples. Results showed that the rice root microbiome of fertilized and non-fertilized fields was not only taxonomically but also functionally distinct (Figure 7). The change in root microbiome function was also observed along with the host plant development stages dependent on the soil fertilization condition except in the seed maturation phase of non-fertilized soil (Figure 7a).

**Figure 7.**
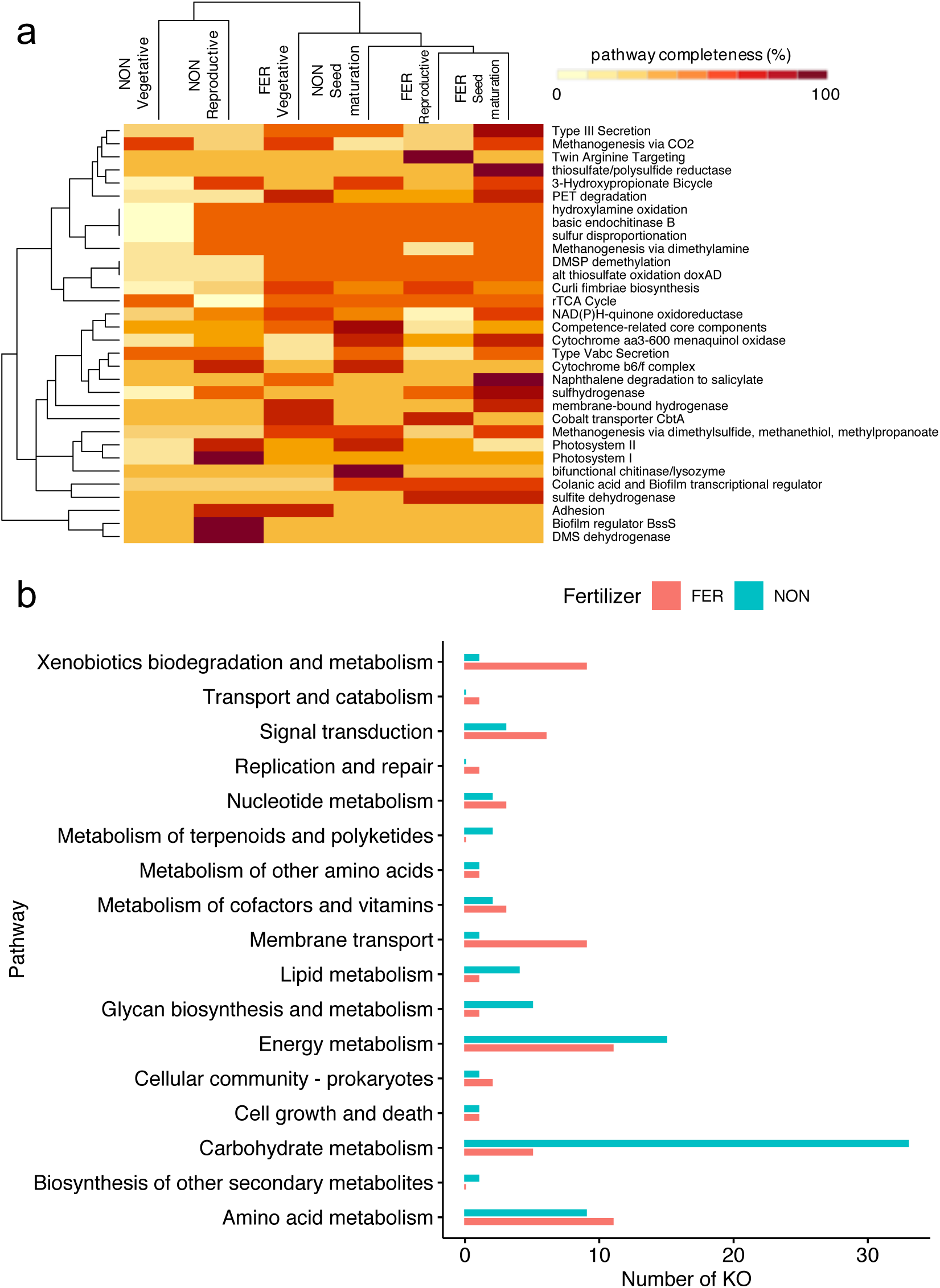
Bacterial functions represented by the root endosphere microbiome of rice in fertilized and non-fertilized fields. (a) Heatmap plot illustrating the completeness of various metabolic pathways based on KEGG (Kyoto Encyclopedia of Genes and Genomes) analysis. The plot highlights pathways with significant differences in completeness between samples from fertilized and non-fertilized fields. (b) Bar plot categorizing relevant KEGG orthologs (KOs) identified by the RF method, showing the distribution of these pathways in both fertilized and non-fertilized fields.

Using a random forest algorithm, we detected that 156 out of 9,304 KOs have a significant difference in relative abundance between non-fertilized (NON) and fertilized (FER) fields. Seventy-three KOs enriched in the non-fertilized field can be classified into 54 pathways, while the rest of them are fertilized field-enriched and can be classified into 47 pathways. Notably, one KO can be assigned to multiple pathways. Interestingly, the proportion of non-fertilized field-enriched KOs related to carbon, lipid, and glycan metabolism was higher than those of fertilized field-enriched KOs (Figure 7b). These three organic compounds are likely to be found on root exudates (Sasse *et al*., 2018). This may indicate that under low nutrition status in the non-fertilized field, plants have higher activity in recruiting microbes via root chemicals to assist them in nutrition provision. In contrast, in fertilized field-enriched KOs, the proportion of KOs related to xenobiotics biodegradation and metabolism are higher than non-fertilized field. Xenobiotic compounds are usually contained in fertilizers and pesticides (Prashar & Shah, 2016). Indeed, this indicates that the application of synthetic chemicals during the farming process in conventional rice fields directly affects the root microbiome metabolism.

## Discussion

### Rice root microbiomes are influenced by soil fertilization conditions and plant developmental stage

Our analyses on a total of 399 root endosphere microbiomes (Table S2) from rice grown in neighboring paddy fields with nutrient-poor (non-fertilized) and nutrient-rich (fertilized field) soils (Table S1), suggest that the structure of rice root-associated microbiomes is primarily influenced by soil fertilization status and plant developmental stage (Figures 2 and 5), with a lesser impact from host genotype (Table S3 and S6). While the effect of fertilization on alpha-diversity (species richness) shows some variability, its impact on beta-diversity (species turnover) aligns with previous studies (Ikeda *et al*., 2014; Sinong *et al*., 2021). Our RF model indicates that differences in root microbiomes between fertilized and non-fertilized soils are most pronounced at approximately 13-19 weeks after germination, around the vegetative-to-reproductive phase transition (Figure 6c). The developmental stage-dependent shifts in the root microbiome are partly correlated with increased alpha diversity, as shown by the Shannon index analyses (Figure 1b). This variation may be due to changes in root exudate composition, which evolves with plant development (Dondjou *et al*., 2023) and affects microbiome composition (Pascale *et al*., 2020). Compared to soil fertilization and plant development stage, differences in microbiome diversity among the three Japonica rice genotypes (Nipponbare, Hinohikari, and Kinmaze) are less pronounced, consistent with studies on other Japonica varieties in controlled conditions (Edwards *et al*., 2015; Sinong *et al*., 2021). Overall, this study demonstrates that soil fertilization and plant age interact to shape the rice root microbiome, which continuously shifts throughout the plant’s life cycle while remaining distinct based on soil fertilization status.

Our data suggest that the impact of *CCaMK* disruption on microbiome assembly is influenced by environmental conditions. Ikeda *et al*. (2011) found minimal changes in bacterial associations in laboratory-grown rice with *ccamk* or other common symbiosis pathway (CSP) mutations. However, in field-growth rice, *CCaMK* affects root endosymbiotic microbiome diversity and plant growth. Our study showed no significant difference in alpha diversity between WT and *ccamk* roots in either fertilized or non-fertilized fields. Conversely, beta diversity differed significantly between WT and *ccamk* only under greenhouse conditions. Notably, the divergent root microbiomes of WT and *ccamk* tend to converge at latter developmental stages (Figure 5f), aligning with Edwards *et al*. (2018), suggesting that *CCaMK* mainly influences initial colonizers in greenhouse. In paddy fields, however, *CCaMK* disruption did not significantly impact root microbiome assembly, suggesting that other dominant factors at play. Factors such as lower cation exchange capacity (Table S1) and different initial soil microbiome (Figure S2a and S2b) may contribute to these field results. Overall, our findings support the notion that CCaMK effects on the root microbiome is environmentally dependent. While CCaMK is known to enhance rice tolerance to drought and oxidative stress (Chen *et al*., 2021), its role in microbiome establishment varies with environmental conditions, highlighting its condition-dependent function and the potential for CCaMK-independent beneficial interactions in paddy rice.

### The root endosphere microbiome of rice grown in non-fertilized field is adaptive to nutrient-deficient soils

After confirming the impact of fertilization conditions on root microbiome assembly, we identified specific microbes contributing to the differences between fertilized and non-fertilized fields. Using ALDEx2, our analyses on relative microbial abundance identified 183 significant ASVs (Figure 3), with 61 over-represented in the non-fertilized field (Table S4). Notably, 30% of these ASVs were assigned to nitrogen-fixing genera like *Rhizomicrobium*, *Azospirillum*, and *Bradyrhizobium*. This over-representation was confirmed by a RF model, which highlighted *Telmatospirillum*, *Bradyrhizobium,* and *Rhizomicrobium* as key indicators for non-fertilized soils (Figure 6b). This may explain the consistent nitrogen uptake despite different fertilization regimes (Figure S4). Additionally, bacteria involved in ferric iron reduction, such as *Telmatospirillum* (Gagen *et al*., 2019), *Uliginosibacterium* (Han *et al*., 2018), and *Geothrix* (Han *et al*., 2023), were abundant in non-fertilized field roots, possibly enhancing iron uptake (Figure S3a). The acidic conditions in non-fertilized field soils likely contribute to increased iron and manganese availability (Table S1; Rengel, 2015). Metagenome analysis revealed distinct functions of microbiomes formed in fertilized versus non-fertilized soils (Figure 7a). While no specific pathways for nutrient acquisition were identified, KO categorization showed that non-fertilized field microbiomes focus on macronutrient metabolism (Figure 7b), implying higher exudation activity from plants to attract beneficial microbes, like *Azospirillum* (Table S4; Upadhyay *et al*., 2022). In contrast, the fertilized field showed an overrepresentation of KOs related to xenobiotic metabolism, indicating that anthropogenic chemicals in conventional farming may influence microbial communities.

### Prediction of soil fertilization status using machine learning models based on plant root microbiomes

Our studies show that rice root microbiomes can effectively predict soil nutritional conditions. Microbiomes integrate both abiotic and biotic factors, making them valuable for forecasting ecosystem processes (Correa-Garcia *et al*., 2023). Recent research has utilized machine learning to predict soil health and crop productivity from soil microbiome profiles (Yergeau *et al*., 2020; Wilhelm *et al*., 2022). While many studies focus on soil microbiomes, the predictive potential of plant microbiomes remains less clear due to their interactions with plant roots. Some studies have used machine learning to identify bacterial biomarkers in rice microbiomes, mostly for classifying rice cultivars or disease resistance (Zhang *et al*., 2019; Xiong *et al*., 2021; Cheng *et al*., 2023; Emmenegger *et al*., 2023). Our study, however, has successfully predicted soil fertilization states using microbiome profiles and identified potential bacterial biomarkers for distinguishing between fertilization conditions.

To enhance practical application, it is crucial to define the model’s predictive range and link plant root microbiomes to soil micronutrient contents. Additionally, like predicting wheat grain quality from the soil microbiomes (Asad *et al*., 2023), sampling timing is vital. Our findings, illustrated in Figures 2b and 5e, reveal that the rice microbiome at 13–19 weeks post-germination is most effective for classifying soil nutrition conditions, indicating stable associations between root-colonizing microbes and soil nutrients during this period. This research advances our understanding of plant-microbe-soil interactions and underscores the practical utility of using rice root microbiomes to assess soil nutrition.

## Acknowledgements

We thank NPO corporation “No Organic or Chemical Input Crop Production Research Group” (NPO *Muhiken*, Kyoto, Japan) for their management and permission to use the rice fields, and Yukiko Shimizu, Yusa Aritoshi, Shota Kido, Miki Yokote, and Maki Saiki for technical assistance. This work was supported in part by the grants from the Canon Foundation “Pursuit of Ideals”, MEXT of Japan 18H02467 (YS), and JST SPRING, Japan Grant Number JPMJSP2140 (AA).

## Author contributions

YS conceived the study. AA, YDU and YS designed the experiments. AA, YDU, DJJA, MF, SK, SI, TM, RS, TK and TF performed the experiments and analyzed the data. KM developed the materials. TM, YH, NO and SK advised on the data analyses. AA, YDU, DJJA and YS wrote the manuscript with contributions from the other authors.

## Data availability

The 16S rRNA gene amplicon and metagenome sequencing data have been deposited with links to BioProject accession number PRJDB18681 and PRJDB18727, respectively, in the DDBJ BioProject database.

## Supporting Information

**Figure S1.**
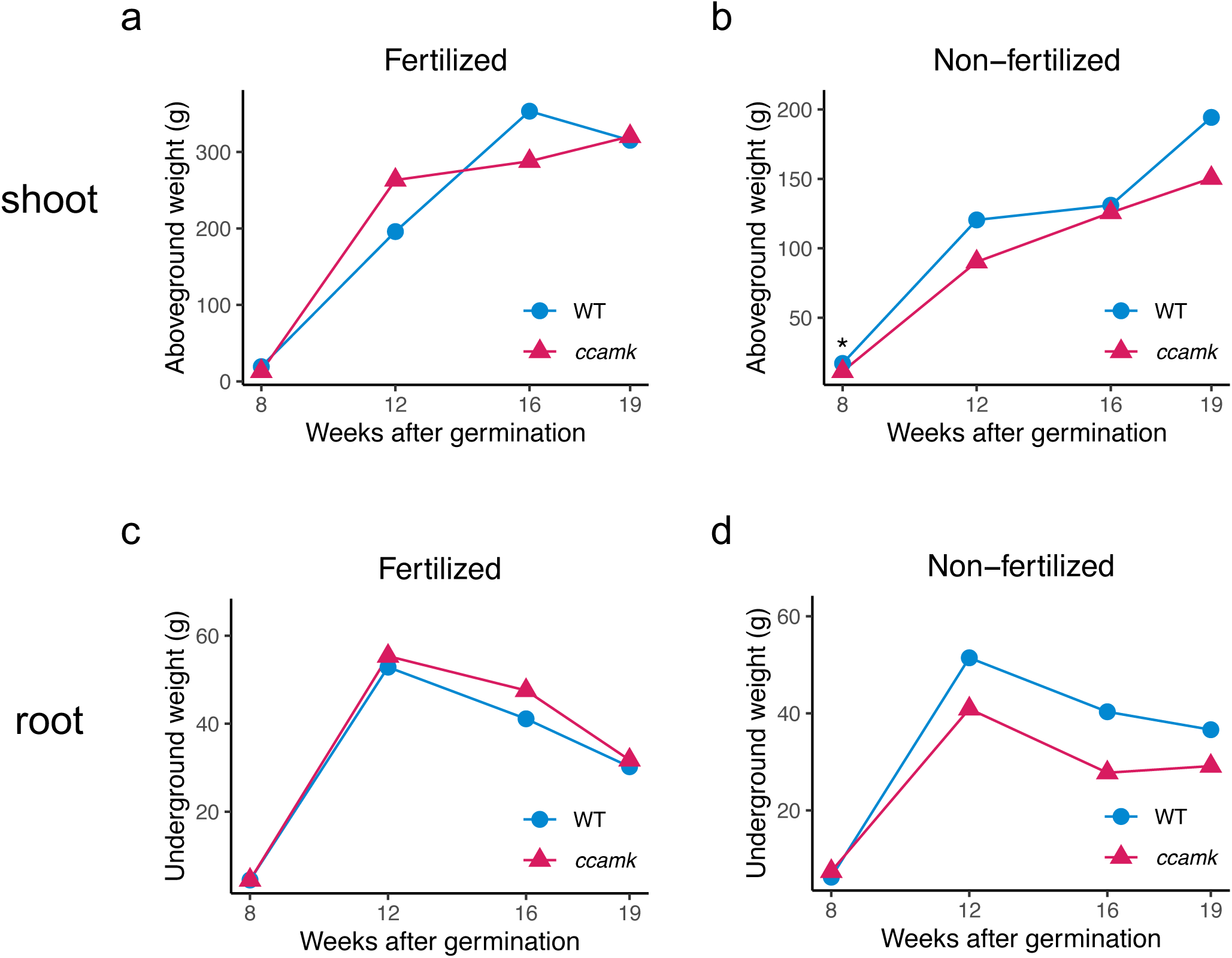
Aboveground and underground weights of wild-type (WT) Nipponbare and *ccamk* mutants. Changes in the rice plant weight of WT and *ccamk* mutants from 8 to 19 weeks after germination in the fields in 2021. The aboveground weights under fertilized and non-fertilized conditions are shown in (a) and (b), respectively. The underground weights under fertilized and non-fertilized conditions are shown in (c) and (d), respectively. Two-sided unpaired Welch’s t-tests were performed with Bonferroni correction applied. Asterisks indicate significant differences (*: adjusted *p* < 0.05).

**Figure S2.**
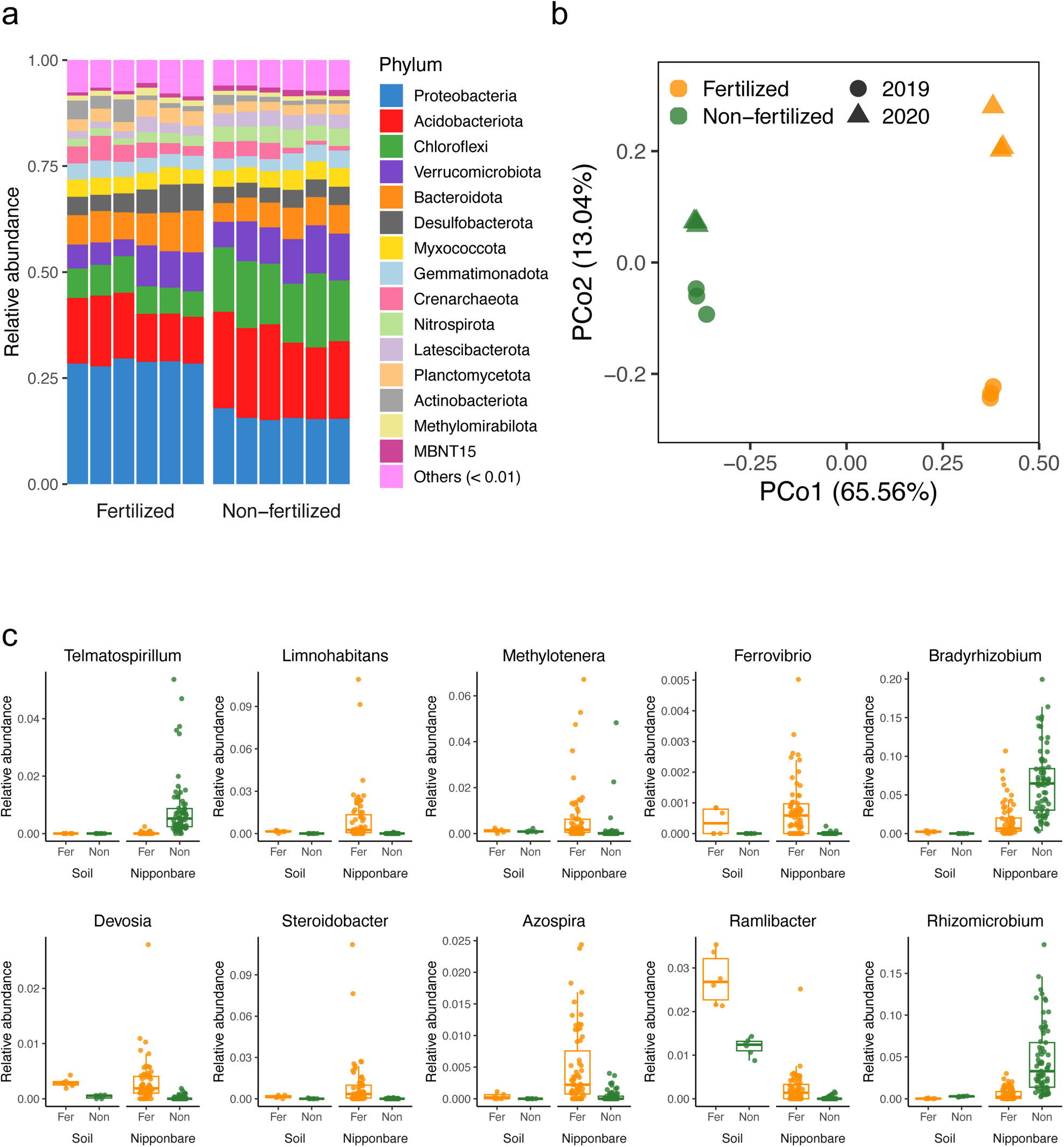
Microbiome profiling of bulk soil from the fertilized and non-fertilized field. a) Phylum level relative abundance of ASVs in the soil microbiome. b) The principal coordinate analysis (PCoA) plots of Bray-Curtis dissimilarities between soil microbiome samples, colored by soil fertilization conditions. c) Comparison of candidate biomarkers’ abundance in the soil versus roots, differentiated by soil fertilization conditions. These biomarkers, selected based on their importance in predicting soil nutrient conditions from a trained random forest model, were assessed in samples collected four weeks after germination in 2019 and 2020. Fer, fertilized field; Non, non-fertilized field.

**Figure S3.**
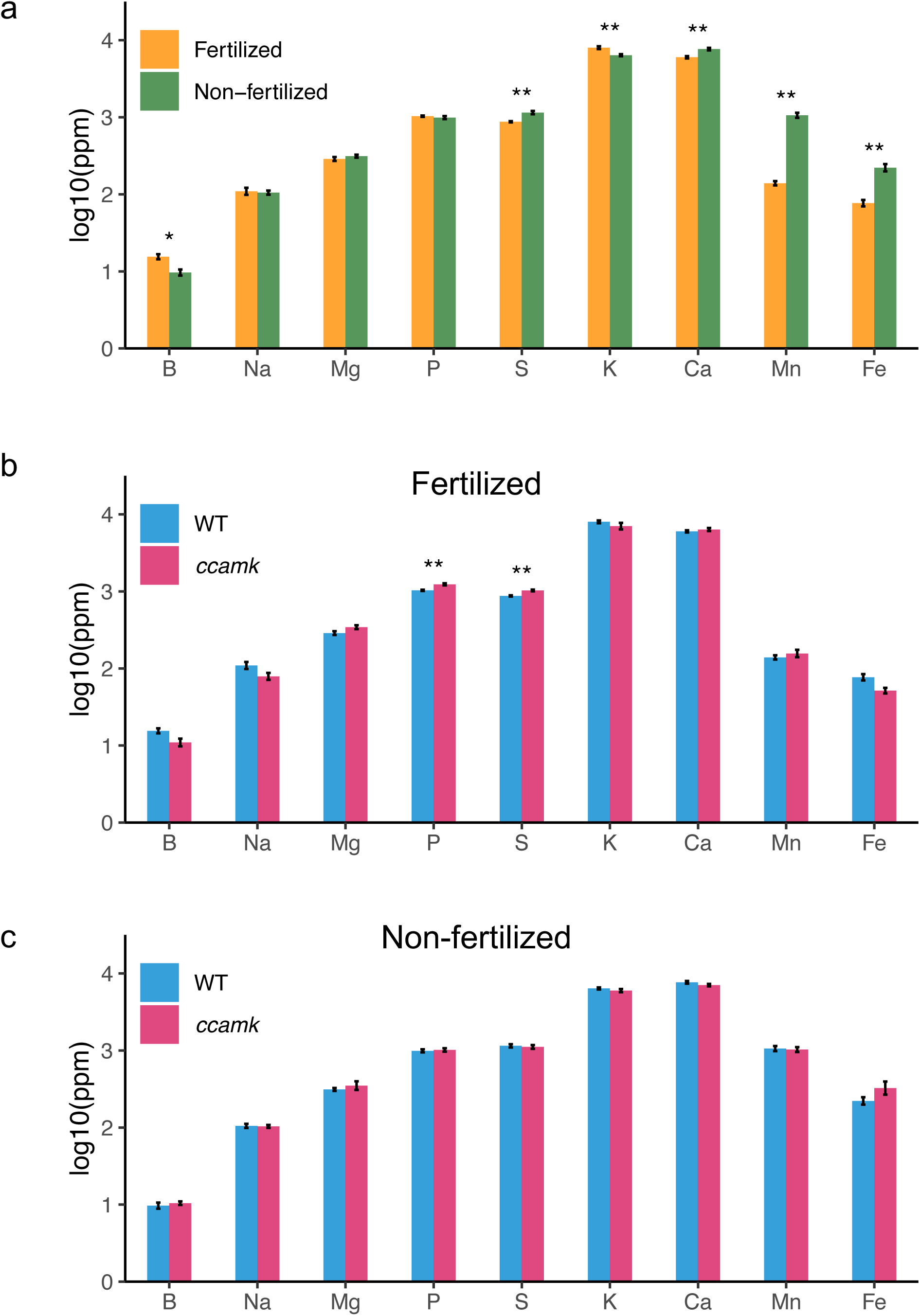
Comparison of elements in the stover leaves of rice grown in the fertilized and non-fertilized field. ICP-MS analysis of elements in stover leaves of (a) wild-type rice grown in the fertilized (n = 10) and non-fertilized fields (n = 10), and wild-type rice and *ccamk* mutant grown in fertilized (n = 10 and n = 9, respectively) (b) and non-fertilized field (n = 10 for both) (c) in 2021. Error bars represent standard errors of the mean. Two-sided unpaired Welch’s t-tests were performed, and Bonferroni correction was applied. Asterisks indicate significant differences (*: adjusted *p* < 0.05, **: adjusted *p* < 0.01).

**Figure S4.**
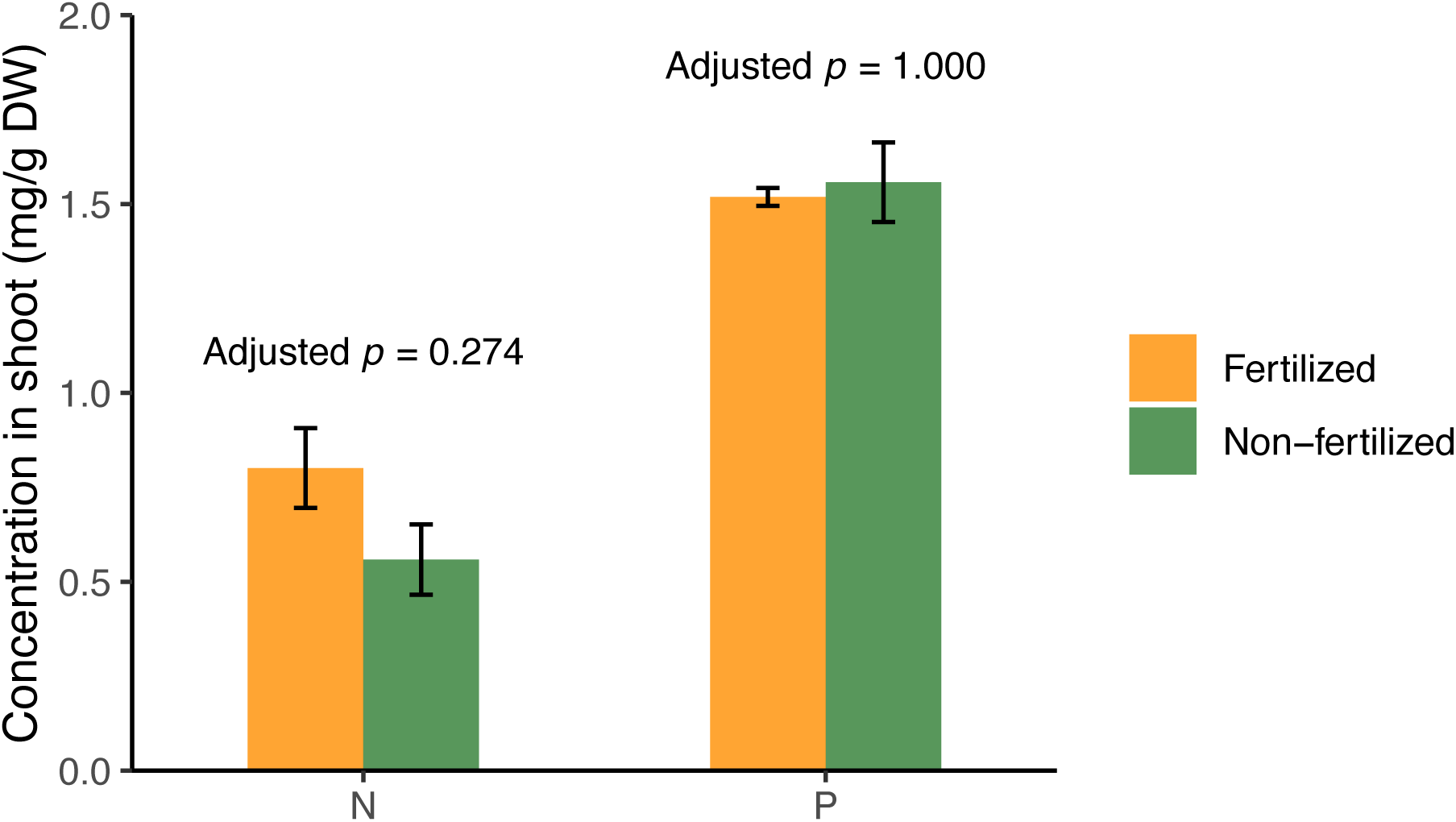
Comparison of nitrogen (N) and phosphorus (P) uptake in rice plants grown in fertilized and non-fertilized fields. Nitrogen (N) and phosphorus (P) concentrations were measured in rice shoots of Nipponbare 18 weeks after germination in 2019. The data were compared between fertilized (n = 4) and non-fertilized fields (n = 4). Error bars represent standard errors of the mean. Two-sided unpaired Welch’s t-tests were performed, with Bonferroni correction applied. No significant differences were observed between the conditions. DW, dry weight.

**Table S1** Soil properties, nutrients and minerals of non-fertilized and fertilized soil.

**Table S2** Field sample collection from 2018-2021.

**Table S3** PERMANOVA to determine the significance of variables in the root microbiome assembly of different rice cultivars in the field.

**Table S4** List of ASVs over-represented in the non-fertilized field.

**Table S5** Root samples collected from wild-type and *ccamk* rice plants grown in the greenhouse.

**Table S6** PERMANOVA to determine the significance of variables in the root microbiome assembly of the wild type and *ccamk* plants grown in the field.

**Table S7** PERMANOVA to determine the significance of variables in the root microbiome assembly of wild type and *ccamk* mutant grown in the greenhouse.

