## Supplementary Table for "Soil nutrition-dependent dynamics of the root-associated microbiome in paddy rice"

**Table S1. Soil properties, nutrients and minerals of non-fertilized and fertilized soil.**

|  | Non-fertilized | | | | Fertilized | | | |
| --- | --- | --- | --- | --- | --- | --- | --- | --- |
|  | actual value | lower limit | upper limit | unit | actual value | lower limit | upper limit | unit |
| pH | 5.3 | 5.4 | 6.4 | - | 6.7 | 5.4 | 6.4 | - |
| EC | 0.02 | - | 0.3 | mS/cm | 0.08 | - | 0.3 | mS/cm |
| Humus | 1.8 | 3 | - | % | 3.7 | 3 | - | % |
| CEC | 14 | 12 | - | meq/100g dry soil | 18 | 12 | - | meq/100g dry soil |
| Ammonia | 1.9 | - | - | mg/100g | 2.4 | - | - | mg/100g |
| Nitrate | 0.1 | - | - |  | - | - | - |  |
| Inorganic nitrogen | 2 | - | - |  | 2.4 | - | - |  |
| Organic phosphate | 4 | 10 | 50 |  | 97 | 10 | 50 |  |
| Potassium | 6 | 14 | 68 |  | 23 | 14 | 68 |  |
| Calcium | 148 | 263 | 304 |  | 471 | 263 | 304 |  |
| Magnesium | 19 | 58 | 72 |  | 20 | 58 | 72 |  |
| Boron | 0.2 | 0.5 | 2 | mg/kg | 0.4 | 0.5 | 2 | mg/kg |
| Manganese | 67.1 | 5 | 8 |  | 6.1 | 5 | 8 |  |
| Zinc | 5.5 | 8 | 40 |  | 26.2 | 8 | 40 |  |
| Copper | 11.4 | 0.8 | 2 |  | 1.5 | 0.8 | 2 |  |
| Silicic acid | 14.8 | 15 | - | mg/100g | 29.8 | 15 | - | mg/100g |

*Blue indicates below the standard while red indicates above the standard

**Table S2. Field sample collection from 2018-2021.**

| **Variety** | **Sampling year** | **Sampling week** | **#Fer** | **#Non** |
| --- | --- | --- | --- | --- |
| Nipponbare | 2018 | 4, 6, 8, 10, 12, 14, 16, 18, 20 | 27 | 24 |
|  | 2019 | 4, 6, 8, 12, 15, 18 | 18 | 18 |
|  | 2020 | 4, 6, 8, 10, 12, 14, 16, 20, 22 | 18 | 27 |
|  | 2021 | 5, 8, 12, 15, 19 | 15 | 15 |
| Hinohikari | 2019 | 4, 6, 8, 12, 15, 18 | 18 | 18 |
|  | 2020 | 4, 6, 8, 10, 12, 14, 16, 20, 22 | 27 | 27 |
| Kinmaze | 2018 | 6, 8, 10, 12, 14, 16, 18, 20 | 24 | 24 |
| *ccamk* | 2019 | 4, 15, 18 | 9 | 9 |
|  | 2020 | 4, 6, 8, 10, 12, 14, 16, 20, 22 | 24 | 27 |
|  | 2021 | 5, 8, 12, 15, 19 | 15 | 15 |
| Soil | 2019 | 4 | 3 | 3 |
|  | 2020 | 4 | 3 | 3 |

*#Fer and #Non means the number of samples in the fertilized and non-fertilized fields, respectively.

**Table S3. PERMANOVA to determine the significance of variables in the root microbiome assembly of different rice cultivars in the field.**

|  | **Df** | **SumOfSqs** | **R^2^** | **F** | **Pr(>F)** |
| --- | --- | --- | --- | --- | --- |
| Fertilized condition | 1 | 7.762 | 0.07376 | 24.6858 | 0.001 |
| Genotype | 2 | 3.905 | 0.03711 | 6.2101 | 0.001 |
| Fertilized condition: Genotype | 2 | 1.125 | 0.01069 | 1.7888 | 0.001 |
| Residual | 294 | 92.441 | 0.87844 |  |  |
| Total | 299 | 105.233 | 1.00000 |  |  |

**Table S4. List of ASVs over-represented in the non-fertilized field.**

| ASV | Kingdom | Phylum | Class | Order | Family | Genus | Species |
| --- | --- | --- | --- | --- | --- | --- | --- |
| ASV2 | Bacteria | Proteobacteria | Alphaproteobacteria | Rhizobiales | Xanthobacteraceae | Bradyrhizobium | Unclassified |
| ASV8 | Bacteria | Proteobacteria | Gammaproteobacteria | Burkholderiales | Burkholderiaceae | Burkholderia-Caballeronia-Paraburkholderia | Unclassified |
| ASV11 | Bacteria | Proteobacteria | Gammaproteobacteria | Burkholderiales | Rhodocyclaceae | Uliginosibacterium | Unclassified |
| ASV17 | Bacteria | Actinobacteriota | Actinobacteria | Streptomycetales | Streptomycetaceae | Streptomyces | Unclassified |
| ASV29 | Bacteria | Proteobacteria | Alphaproteobacteria | Rhizobiales | Xanthobacteraceae | Unclassified | Unclassified |
| ASV32 | Bacteria | Proteobacteria | Gammaproteobacteria | Burkholderiales | Burkholderiaceae | Ralstonia | Unclassified |
| ASV34 | Bacteria | Actinobacteriota | Actinobacteria | Kineosporiales | Kineosporiaceae | Unclassified | Unclassified |
| ASV38 | Bacteria | Proteobacteria | Alphaproteobacteria | Micropepsales | Micropepsaceae | Rhizomicrobium | Unclassified |
| ASV45 | Bacteria | Proteobacteria | Gammaproteobacteria | Burkholderiales | Comamonadaceae | Ideonella | dechloratans |
| ASV58 | Bacteria | Proteobacteria | Alphaproteobacteria | Micropepsales | Micropepsaceae | Rhizomicrobium | electricum |
| ASV67 | Bacteria | Proteobacteria | Gammaproteobacteria | Burkholderiales | Rhodocyclaceae | Denitratisoma | Unclassified |
| ASV70 | Bacteria | Proteobacteria | Gammaproteobacteria | Burkholderiales | Oxalobacteraceae | Herbaspirillum | Unclassified |
| ASV88 | Bacteria | Proteobacteria | Alphaproteobacteria | Rhizobiales | Beijerinckiaceae | Methylosinus | trichosporium |
| ASV89 | Bacteria | Proteobacteria | Alphaproteobacteria | Rhizobiales | Pleomorphomonadaceae | Unclassified | Unclassified |
| ASV96 | Bacteria | Proteobacteria | Alphaproteobacteria | Azospirillales | Azospirillaceae | Azospirillum | Unclassified |
| ASV99 | Bacteria | Acidobacteriota | Holophagae | Holophagales | Holophagaceae | Unclassified | Unclassified |
| ASV106 | Bacteria | Proteobacteria | Gammaproteobacteria | Burkholderiales | Rhodocyclaceae | Uliginosibacterium | Unclassified |
| ASV107 | Bacteria | Proteobacteria | Alphaproteobacteria | Micropepsales | Micropepsaceae | Rhizomicrobium | Unclassified |
| ASV111 | Bacteria | Acidobacteriota | Holophagae | Holophagales | Holophagaceae | Geothrix | Unclassified |
| ASV116 | Bacteria | Acidobacteriota | Acidobacteriae | Acidobacteriales | Acidobacteriaceae (Subgroup 1) | Unclassified | Unclassified |
| ASV120 | Bacteria | Proteobacteria | Alphaproteobacteria | Rhizobiales | Pleomorphomonadaceae | Unclassified | Unclassified |
| ASV130 | Bacteria | Proteobacteria | Alphaproteobacteria | Micropepsales | Micropepsaceae | Rhizomicrobium | Unclassified |
| ASV133 | Bacteria | Proteobacteria | Alphaproteobacteria | Rhizobiales | Xanthobacteraceae | Bradyrhizobium | Unclassified |
| ASV137 | Bacteria | Spirochaetota | Spirochaetia | Spirochaetales | Spirochaetaceae | Salinispira | Unclassified |
| ASV143 | Bacteria | Proteobacteria | Alphaproteobacteria | Azospirillales | Azospirillaceae | Azospirillum | Unclassified |
| ASV145 | Bacteria | Proteobacteria | Gammaproteobacteria | Burkholderiales | Comamonadaceae | Curvibacter | Unclassified |
| ASV148 | Bacteria | Proteobacteria | Alphaproteobacteria | Rhodospirillales | Magnetospirillaceae | Telmatospirillum | Unclassified |
| ASV154 | Bacteria | Proteobacteria | Gammaproteobacteria | Burkholderiales | Rhodocyclaceae | Propionivibrio | Unclassified |
| ASV157 | Bacteria | Bacteroidota | Bacteroidia | Chitinophagales | Chitinophagaceae | Chitinophaga | oryziterrae |
| ASV160 | Bacteria | Firmicutes | Clostridia | Oscillospirales | Oscillospiraceae | Sporobacter | Unclassified |
| ASV183 | Bacteria | Proteobacteria | Gammaproteobacteria | Pseudomonadales | Moraxellaceae | Acinetobacter | Unclassified |
| ASV188 | Bacteria | Proteobacteria | Alphaproteobacteria | Micropepsales | Micropepsaceae | Rhizomicrobium | Unclassified |
| ASV195 | Bacteria | Acidobacteriota | Holophagae | Holophagales | Holophagaceae | Unclassified | Unclassified |
| ASV199 | Bacteria | Proteobacteria | Alphaproteobacteria | Azospirillales | Azospirillaceae | Azospirillum | Unclassified |
| ASV224 | Bacteria | Proteobacteria | Alphaproteobacteria | Rhodospirillales | Magnetospirillaceae | Telmatospirillum | Unclassified |
| ASV229 | Bacteria | Bacteroidota | Bacteroidia | Chitinophagales | Chitinophagaceae | Niastella | Unclassified |
| ASV239 | Bacteria | Firmicutes | Negativicutes | Veillonellales-Selenomonadales | Sporomusaceae | Unclassified | Unclassified |
| ASV258 | Bacteria | Acidobacteriota | Holophagae | Holophagales | Holophagaceae | Unclassified | Unclassified |
| ASV285 | Bacteria | Acidobacteriota | Acidobacteriae | Acidobacteriales | Acidobacteriaceae (Subgroup 1) | Terracidiphilus | Unclassified |
| ASV291 | Bacteria | Proteobacteria | Alphaproteobacteria | Azospirillales | Azospirillaceae | Azospirillum | Unclassified |
| ASV306 | Bacteria | Desulfobacterota | Desulfovibrionia | Desulfovibrionales | Desulfovibrionaceae | Desulfovibrio | Unclassified |
| ASV310 | Bacteria | Patescibacteria | Saccharimonadia | Saccharimonadales | Unclassified | Unclassified | Unclassified |
| ASV314 | Bacteria | Proteobacteria | Gammaproteobacteria | Gammaproteobacteria Incertae Sedis | Unknown Family | Acidibacter | Unclassified |
| ASV326 | Bacteria | Proteobacteria | Alphaproteobacteria | Micropepsales | Micropepsaceae | Rhizomicrobium | Unclassified |
| ASV344 | Bacteria | Proteobacteria | Alphaproteobacteria | Azospirillales | Azospirillaceae | Azospirillum | Unclassified |
| ASV358 | Bacteria | Acidobacteriota | Acidobacteriae | Acidobacteriales | Acidobacteriaceae (Subgroup 1) | Terracidiphilus | Unclassified |
| ASV360 | Bacteria | Proteobacteria | Gammaproteobacteria | Pseudomonadales | Pseudomonadaceae | Pseudomonas | Unclassified |
| ASV392 | Bacteria | Bacteroidota | Bacteroidia | Chitinophagales | Chitinophagaceae | Niastella | vici |
| ASV405 | Bacteria | Firmicutes | Negativicutes | Veillonellales-Selenomonadales | Sporomusaceae | Unclassified | Unclassified |
| ASV420 | Bacteria | Proteobacteria | Alphaproteobacteria | Dongiales | Dongiaceae | Dongia | Unclassified |
| ASV424 | Bacteria | Myxococcota | Polyangia | Haliangiales | Haliangiaceae | Haliangium | Unclassified |
| ASV462 | Bacteria | Verrucomicrobiota | Verrucomicrobiae | Opitutales | Opitutaceae | Opitutus | Unclassified |
| ASV581 | Bacteria | Spirochaetota | Spirochaetia | Spirochaetales | Spirochaetaceae | Unclassified | Unclassified |
| ASV584 | Bacteria | Spirochaetota | Spirochaetia | Spirochaetales | Spirochaetaceae | Salinispira | Unclassified |
| ASV617 | Bacteria | Proteobacteria | Alphaproteobacteria | Micropepsales | Micropepsaceae | Rhizomicrobium | Unclassified |
| ASV679 | Bacteria | Armatimonadota | Fimbriimonadia | Fimbriimonadales | Fimbriimonadaceae | Unclassified | Unclassified |
| ASV690 | Bacteria | Bacteroidota | Bacteroidia | Bacteroidales | Bacteroidetes vadinHA17 | Unclassified | Unclassified |
| ASV722 | Bacteria | Bacteroidota | Bacteroidia | Bacteroidales | Bacteroidetes BD2-2 | Unclassified | Unclassified |
| ASV749 | Bacteria | Proteobacteria | Alphaproteobacteria | Acetobacterales | Unclassified | Unclassified | Unclassified |
| ASV831 | Bacteria | Myxococcota | Myxococcia | Myxococcales | Anaeromyxobacteraceae | Anaeromyxobacter | Unclassified |
| ASV994 | Bacteria | Spirochaetota | Spirochaetia | Spirochaetales | Spirochaetaceae | Unclassified | Unclassified |

*Green and red indicate possible involvement of ASVs in nitrogen fixation and iron reduction, respectively.

**Table S5. Root samples collected from wild-type and *ccamk* rice plants grown in the greenhouse.**

| **Variety** | **Sampling year** | **Sampling week** | **#Fer** | **#Non** |
| --- | --- | --- | --- | --- |
| Nipponbare | 2020 | 6, 10, 12, 17 | 16 | 16 |
| *ccamk* | 2020 | 6, 10, 12, 17 | 16 | 16 |

*#Fer and #Non represents the total number of samples (4 replicates per time point) in the fertilized and non-fertilized pots, respectively.

**Table S6. PERMANOVA to determine the significance of variables in the root microbiome assembly of the wild type and *ccamk* plants grown in the field.**

|  | **Df** | **SumOfSqs** | **R^2^** | **F** | **Pr(>F)** |
| --- | --- | --- | --- | --- | --- |
| Fertilized condition | 1 | 6.732 | 0.09723 | 22.4392 | 0.001 |
| Genotype | 1 | 0.366 | 0.00529 | 1.2211 | 0.204 |
| Fertilized condition:Genotype | 1 | 0.336 | 0.00485 | 1.1185 | 0.293 |
| Residual | 206 | 61.799 | 0.89263 |  |  |
| Total | 209 | 69.233 | 1.00000 |  |  |

**Table S7. PERMANOVA to determine the significance of variables in the root microbiome assembly of wild type and *ccamk* mutant grown in the greenhouse.**

|  | **Df** | **SumOfSqs** | **R^2^** | **F** | **Pr(>F)** |
| --- | --- | --- | --- | --- | --- |
| Fertilized condition | 1 | 5.1348 | 0.28944 | 28.750 | 0.001 |
| Genotype | 1 | 1.1443 | 0.06450 | 6.407 | 0.001 |
| Fertilized condition:Genotype | 1 | 0.7448 | 0.04198 | 4.170 | 0.002 |
| Residual | 60 | 10.7162 | 0.60407 |  |  |
| Total | 63 | 17.7401 | 1.00000 |  |  |
